# Protocols to enable fluorescence microscopy of microbial interactions on living maize silks (style tissue) that balance the requirements for containment, biological relevance, viability, and experimental accessibility

**DOI:** 10.1101/2023.10.31.564993

**Authors:** M.E.H Thompson, M.N. Raizada

**Affiliations:** Department of Plant Agriculture, University of Guelph, Guelph, ON, Canada

**Keywords:** maize, confocal, fluorescence, biocontrol, *Fusarium*, *F. graminearum*, mycotoxin, beneficial microbe, silk, pollen, pollen tube, fertilization, microbiome, reproduction, pathogen

## Abstract

There is interest in studying microbes that colonize maize silks (style tissue, critical for reproduction) including the fungal pathogen *Fusarium graminearum* (*Fg*) and its interactions with the microbiome and biocontrol agents. *In planta* imaging of these interactions on living silks using confocal fluorescence microscopy would provide key insights. However, newly discovered microbes have unknown effects of on human health, and there are regulatory requirements to prevent the release of fluorescently-tagged microbes into the environment.

Therefore, the microbe infection, colonization, and interaction stages on silks prior to microscopy must be contained. At the same time, silk viability must be maintained and experiments conducted that are biologically relevant (e.g. silks should remain attached to the cob), yet the silk tissue must be accessible to the researcher (i.e. not within husk leaves) and allow for multiple replicates. Here we present methods that meet these five contrasting criteria. We tested these methods using *Fg* and four silk-derived bacterial endophytes. The endophytes were previously known to have anti-*Fg* activity *in vitro*, but *in planta* observations were lacking. In Method 1, a portion of the tip of a cob was dissected, and silks remained attached to the cob in a Petri dish. The cob was placed on a water agar disc to maintain hydration. DsRed-tagged bacteria and GFP-tagged *Fg* were inoculated onto the silks and incubated, allowing the two microbes to grow towards one another before staining with propidium iodide for confocal microscopy. A variation of the protocol was presented in Method 2, where detached silk segments were placed directly on water agar where they were inoculated with bacteria and *Fg* to promote dense colonization, and to allow for many replicates and interventions such as silk wounding. The bacterial endophytes were successfully observed colonizing *Fg* hyphae, silk trichomes, and entering silks via cut ends and wounds. These protocols can be used to study other silk-associated microbes including several globally important fungal pathogens that enter maize grain through silks.

## INTRODUCTION

Plants, including crops, are severely impacted by pathogens. *In planta* imaging can help elucidate the interactions between the host tissue, pathogens, and members of the plant microbiome (Cardinale, 2014; Palmieri et al., 2020).

In maize (*Zea mays* L. ssp. *mays*, corn), silks are the female reproductive tract, more broadly known as the style tissue in other angiosperms, through which male gametes are transmitted to the ovule for fertilization (Kiesselbach, 1999; Sauter, 2009). Silks must be exposed to the environment to receive pollen via wind-pollination, which also makes this tissue susceptible to fungal pathogens (Kiesselbach, 1999). Various fungal pathogens of economic importance use the silks as a route to enter the grain (Thompson & Raizada, 2018). Of these, *Fusarium graminearum* (*Fg*) is a prominent, global mycotoxigenic pathogen of maize that can enter via silks and deposit deoxynivalenol (DON) and other mycotoxins in grain (De Ruyck et al., 2015; Oldenburg et al., 2017; Thompson & Raizada, 2018).

Plant microbiomes are known to contain endophytes, subsurface microbes, that can be anti-fungal and protective to the host (Mousa, Schwan, et al., 2015, 2016; Mousa, Shearer, et al., 2015, 2016). Floral structures have been reported to contain anti-pathogenic endophytes (Aleklett et al., 2014; Cui et al., 2020; Ngugi & Scherm, 2006). Maize silks are of particular recent interest for the discovery and understanding of biocontrol microbes, given their critical role in reproduction and pathogen susceptibility (Diniz, Cota, et al., 2022; Diniz, Figueiredo, et al., 2022; Khalaf et al., 2021).

Fluorescence confocal microscopy is a very useful tool for observing plant-endophyte-pathogen relationships on living tissue including at subsurface planes (Cardinale, 2014; Palmieri et al., 2020). Fluorescence microscopy has previously been used to observe *Fg* tagged with green fluorescent protein (GFP) in maize silks (Miller et al., 2004, 2007), and GFP-tagged bacteria and calcofluor-stained *Fg* on millet roots (Mousa, Shearer, et al. 2016). Endophytes have also been observed on plant tissues via electron microscopy (Sankaranarayanan et al., 2023; War Nongkhlaw & Joshi, 2017).

One challenge with studying microbial interactions using fluorescence microscopy is the use of transgenic microbes, which are regulated in many jurisdictions, for fears of release into the environment (McHughen, 2016). Furthermore, the effects of newly discovered endophytic bacteria on humans are unknown. Given these safety concerns, the microbe infection stage, colonization stage, and microbe-microbe interaction stage on plant tissues prior to microscopy must be contained (e.g. not in a greenhouse or field) when it involves transgenic microbes. It is therefore challenging to study microbes in organs that are large, such as the maize female inflorescence (cob) and its silks which are long. Maize silks represent a novel tissue to understand the interactions between fungal pathogens and bacterial endophytes (Khalaf et al., 2021). Methods are needed to contain microbial-treated silks from the environment, while at the same time, keeping the silks alive (e.g. hydrated) while microbes colonize the silks and form interactions with other microbes of interest (e.g. *Fg*), prior to microscopy. At the same time, it is important to conduct experiments that are biologically relevant. In this case, rather than detached silks, an assay in which the silks remain connected to the ovules on the cob would be ideal. This is particularly true in studies involving *Fg* which colonizes silks directionally; it enters silk tips from the environment, and then migrates towards the silk base, ultimately infecting the ovule (Schaafsma et al., 1997), perhaps guided by a required chemotactic signal. Finally, a protocol is needed that allows for multiple replicates and access to the silks (i.e. without the husk leaves), to control the precise location and timing of microbial inoculations.

Here, two novel protocols are presented that enable the use of confocal fluorescence microscopy on maize silks that balance the requirements for containment, biological relevance, viability, experimental accessibility, and replication. The protocols were validated by imaging the interactions between the pathogen *Fusarium graminearum* and four anti-*Fg* endophytic bacterial isolates.

## METHODS

This experiment tested how the microbes would colonize the silks, and colonize *Fg* on silks, including at deliberate wound sites. There were 2 main modifications of the protocol, specifically: Method 1: silks attached to the cob; and Method 2: silks detached from the cob. The protocols presented for observing host-endophyte-pathogen interactions are flexible and should be adjusted based on the specific microbial strains used. Many variations of application timelines for bacteria and fungi, staining and rinses, and wounding of silks were attempted.

### Source of biological materials

Source of transgenic *Fg* strain: Tagged microbes in viable silk tissue were imaged in the presence and absence of GFP-tagged *Fg*. The ZTE-2A strain of GFP-tagged *Fg* (Miller et al., 2004) was provided by Dr. Rajagopal Subramaniam (AAFC, Ottawa). The GFP-*Fg* was inoculated in 25 mL of potato dextrose broth in a 50 mL Falcon tube and incubated at 25°C and 120 rpm for 3 d before it was used for inoculation.

Source of bacterial strains: Bacterial isolates R22 (*Pantoea ananatis*), R67 (*Pseudomonas*), E04 (*Pantoea*), and L59 (*Delftia*) were cultured from pollinated maize silks at the fertilization stage and found to suppress *Fg in vitro*. Isolates R22, R67, and E04 were transformed with tetracycline (Tet) resistant plasmids pSW002-PpsbA-DsRed-Express2 (DsRed; Addgene, 111257) via electroporation (Supplemental Methods S1). Isolate L59 was transformed with kanamycin (Kan) resistant plasmid, pDSK-GFP (GFP) via electroporation (Supplemental Methods S1).

Source of maize cobs: Sequentially-planted greenhouse-grown and field-grown maize cobs were used. Maize inbreds PHRE1 and LH82 were grown in the Crop Science greenhouse at the University of Guelph in 18.9 L pails of large Turface® clay (Regular Athletics texture, Turface Athletics, USA). Further growth conditions can be found in Supplemental Methods S2. Field plants (PHRE1 inbred) were grown at the University of Guelph Elora Research Station. Cobs were harvested when recently-emerged silks were visible outside of the husk (1-3 cm extended; Figure 1A).

**Figure 1.**
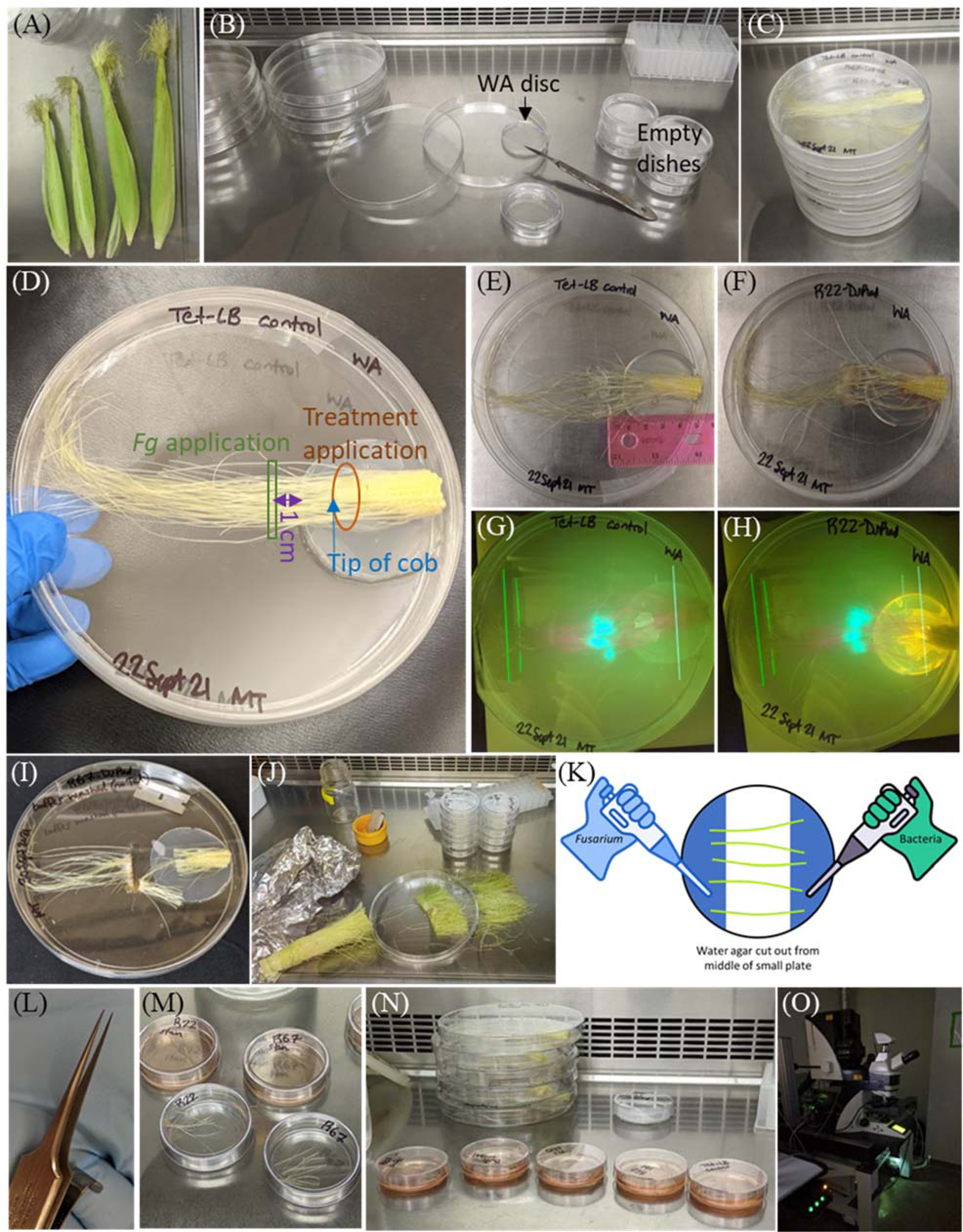
Pre-confocal imaging methods to enable observations of microbial interactions on maize silks using fluorescently-tagged microbes that facilitate containment, biological relevance, viability, experimental accessibility and replication. Shown here was the bacterium *Pantoea ananatis* R22 tagged with DsRed and fungal pathogen *Fusarium graminearum* (*Fg*) tagged with GFP. **(A)** Young maize (corn) cobs with freshly emerged silks are used in these protocols. **(B)** For ***Method 1 - Silks attached to the cob***, Step 1: a small water agar plate (12 g water agar/1 L ddH2O, 10 mL per plate, 60 mm diameter) was removed from the original Petri dish and placed in a 150 mm diameter Petri dish so that the water agar touched one edge of the large dish. **(C)** Step 2: The husks were removed from young cobs with freshly emerged silks and the cobs were cut 3 cm from the tip, and cut again lengthwise with a sterile razor blade. Loose or severed silks were removed, and half of a cob was plated onto the small circle of water agar with the silks extending across the Petri dish. **(D)** Step 3: Applications of 400 µL (in 2 aliquots of 200 µL) of Tris-HCl-washed bacterial liquid culture or 10 mM Tris-HCl control treatment and GFP-tagged *Fg* (either at the same time as the treatment, or 1 d later) were applied onto the designated locations. The GFP-tagged *Fg* was briefly vortexed thrice, until fragments of hyphae broke apart from the main mass, before small pieces of *Fg* were applied using a 1000 µL pipette. The plates were kept level as Petri dishes were sealed with Parafilm and incubated in a dark, 25°C incubator. **(E-H)** After incubation (Step 4), microbial growth can be seen on control-treated silks **(E)** and silks treated with bacterial strain R22 **(F). (G-H)** Control-treated and R22-treated silks under UV light, after incubation. GFP-tagged *Fg* can be seen fluorescing in both. DsRed-tagged R22 can be seen fluorescing in (H). **(I)** Step 5: After incubation, silks were cut with sterile razors; one cut was made where the fungus had been applied, and the second cut was near the tip of the cob, to make silk segments approximately 2.5 cm long. **(J)** For ***Method 2 - Silks detached from the cob***, the silks were cut prior to bacteria/*Fg* application and incubation, and 5 pieces of silks were placed on a small water agar plate. Silks were treated with 300 µL (in 2 doses of 150 µL) of Tris-HCl-washed bacterial liquid culture or 10 mM Tris-HCl control, and 200 µL of GFP- *Fg* applied in the same manner, with a P1000 pipette in 2 doses of 100 µL on the same day, or 1 d after the bacteria. **(K)** For *Method 3, Silk Bridge*, the middle section of water agar from a 60 mm diameter Petri dish was cut and removed, leaving the remaining portions of water agar on two sides. Approximately 2.5 cm segments of fresh silks (as in J) were laid across the open space to form a “bridge.” Then 300 µL of GFP-*Fg* was applied to one section of water agar, and 300 µL of Tris-HCl-washed bacterial liquid culture or 10 mM Tris-HCl control was applied to the other section. The glove/pipette icon used in this diagram was from https://www.flaticon.com/free-icons. **(L)** As a sub-modification of *Method 2*, silks were wounded with very sharp, fine tweezers, shown here. **(M-N)** Silks in 60 mm Petri dishes for staining with dilute propidium iodide for 30 min. **(O)** Interactions between *Fg*, bacterial isolates, and viable maize silk tissue were observed one silk at a time in a 27 mm Nunc glass-bottom Petri dish (Catalog No.150682, Thermo Scientific, USA), covered with ddH2O and a cover slip, under the Leica TCS SP5 confocal laser scanning microscope (DM 6000B upright microscope, Leica, Germany) located in the Molecular and Cellular Imaging Facility at the University of Guelph, Canada.

### Co-culturing *Fg* and bacteria on silks for confocal microscopy: novel protocols

#### Method 1. Silks attached to the cob: Assembling silk plates, inoculating with bacteria, and preparing silks for staining

The first challenge was to miniaturize the natural environment where fungal pathogens and endophytes interact on maize silks, so that the assay could be contained while maintaining biological relevance, silk viability, and ability to mimic natural infection. It was decided to dissect maize cobs into tip segments, leaving the silks intact on the cob, while maintaining moisture with a disc of water agar. A sterile scalpel was used to cut around the edge of a water agar plate (12 g water agar/1 L ddH2O, 10 mL per plate, 60 mm diameter) which was then removed (flipped) from the small Petri dish and placed into a larger 150 mm diameter Petri dish so that it touched one edge of the large Petri dish (Figure 1B). A young cob with freshly emerged silks was harvested, the husks were removed, and an autoclaved razor blade was used to cut the cob, 3 cm from the tip. Cutting was done on a spare, large Petri dish to keep the silks clean. The 3 cm-long tip portion of the cob (with extending silks) was cut in half lengthwise, and any severed or loose silks were removed. Half of a cob segment was laid into a large Petri dish, onto the small circle of water agar, so the severed end of the cob was nearly touching the edge of the Petri dish, and the silks were extended, nearly touching the opposite edge of the Petri dish (Figure 1C).

The next challenge was to establish colonization of the silks with bacteria and pathogen that could be visualized on and inside of silk tissue, *in planta* rather than simply observing the bacteria and *Fg in vitro*. It was decided that fluorescently-tagged microbes would be grown in liquid cultures and applied onto the silks. To avoid having the liquid cultures simply combine and interact *in vitro* on agar or the Petri dish, the microbes were applied with the intention that the endophyte and *Fg* would have to migrate towards one another on the silks; the endophyte was applied on the silks close to the cob, elevated on a water agar disc, whereas *Fg* was applied to the silks on the lower surface of the Petri dish (Figure 1D). Specifically, 400 µL (in 2 aliquots of 200 µL) of Tris-HCl-washed bacterial liquid culture or 10 mM Tris-HCl control treatment were applied onto the silks at the cob tip, using a 1000 µL pipette. The GFP-tagged *Fg* was applied either at the same time as the treatment, or 1 d after. Keeping the plates flat, the Petri dishes were sealed with Parafilm and incubated in a dark, 25°C incubator for 1 d to allow for bacterial colonization. The GFP-tagged *Fg* (described above) was briefly vortexed thrice, until fragments of hyphae could be seen breaking apart from the main mass. Then, 400 µL (in 2 aliquots of 200 µL, using a 1000 µL pipette) of small chunks of GFP-tagged *Fg* were applied in a straight line, perpendicular to the silks, 1 cm from the edge of the water agar disc. The fungal liquid did not come into contact with the water agar (if it did, the plate was disposed of). The plates were again sealed with Parafilm and returned to the 25°C, dark incubator for colonization to continue (Figure 1E-H).

On the day of microscopy, the sections of interest where the endophyte and pathogen interacted on the silk were cut out using sterile razor blades (Figure 1I). One cut was made where the fungus was applied, 1 cm away from the water agar. The second cut was near the tip of the cob, up to 1 cm down the cob.

#### Method 2. Silks detached from the cob: Assembling the silk plates and inoculating with bacteria

The next challenge was to establish dense, consistent colonization of silks, allowing for observation of interactions when microbes were slow to colonize or if they were not highly motile under the incubation conditions, *and* to allow for many replicates from a single cob, *and* to permit additional interventions including wounding. It was decided to utilize cut segments of silks (Figure 1J), placing them on a small water agar (WA) plate, and applying liquid cultures of an endophyte and *Fg* at the same location. For this modified method, the ∼2.5 cm sections of silks were cut, and 5 pieces were plated onto a small water agar plate. As a sub-modification, the silks were wounded with very sharp, fine tweezers (Figure 1L). Subsequently, 300 µL of Tris-HCl-washed bacterial liquid culture or 10 mM Tris-HCl control was applied over the silks, coating them in 2 doses of 150 µL. Next, 200 µL of GFP-*Fg* was applied in the same manner, with a P1000 pipette in 2 doses of 100 µL on the same day, or 1 d later than the anti-*Fg* bacteria.

### Preparing silks for microscopy

Silks were incubated for a total of 2-6 d. The silk segments were transferred to a 60 mm diameter Petri dish using sterile tweezers, for staining (Figure 1M). For staining, 500 µL propidium iodide (PI) solution (1 mg/mL in water) was diluted in 10 mL autoclaved ddH2O. Fresh dilutions were prepared on each day of staining. The stock bottle of PI was kept in the dark at 4°C. The diluted PI was applied to cover the silks entirely, and the silks were weighed down with a cover slip to ensure full coverage for 30 min (Figure 1N). Next, the stain was gently removed with a P1000 pipette. A small amount of stain was left in the dish to prevent the silks from drying out, and the dish was placed into a secondary container during transport to the microscope facility. Stained and unstained/unwashed silks were compared to determine whether microbes were washed away during the staining process. No rinsing was required. At the microscope, a single piece of silk tissue was placed in a 27 mm Nunc glass-bottom Petri dish (Catalog No.150682, Thermo Scientific, USA) and covered with ddH2O and a cover slip so that the cover slip was submerged. The glass-bottom Petri dish was cleaned thoroughly with 70% ethanol and Kimwipes between samples.

### Confocal microscopy settings and process

Interactions between *Fg,* bacterial isolates, and viable maize silk tissue were observed under a Leica TCS SP5 confocal laser scanning microscope (DM 6000B upright microscope, Leica, Germany) at the Molecular and Cellular Imaging Facility at the University of Guelph, Canada (Figure 1O). An argon laser (488 nm), set at a maximum of 30%, was used to excite the GFP and PI. GFP and PI have different emission wavelength profiles, so they were detected separately while using the same laser for excitation. A HeNe laser (543 nm), set around 50%, was used to excite the DsRed. The argon and HeNe lasers were used in sequence, by frame. The smart gain and smart offset were adjusted for each laser. For exploring the tissue and locating the fungus, the argon laser was primarily used; the HeNe laser was intermittently used to check for bacteria while exploring the tissue.

## RESULTS

The methodologies (Figure 1) enabled successful imaging of the interactions between candidate maize silk-derived endophytes and *Fusarium graminearum* (*Fg*) on living maize silks under controlled conditions (Figures 2 and 3). The specific biological observations are noted in each extensive figure legend. Variations of application timelines for the bacteria and fungus, staining, rinsing, and wounding were attempted, and it was found that two modifications were most effective, specifically (1) silks attached to the cob; (2) silks detached from the cob, as detailed below.

**Figure 2.**
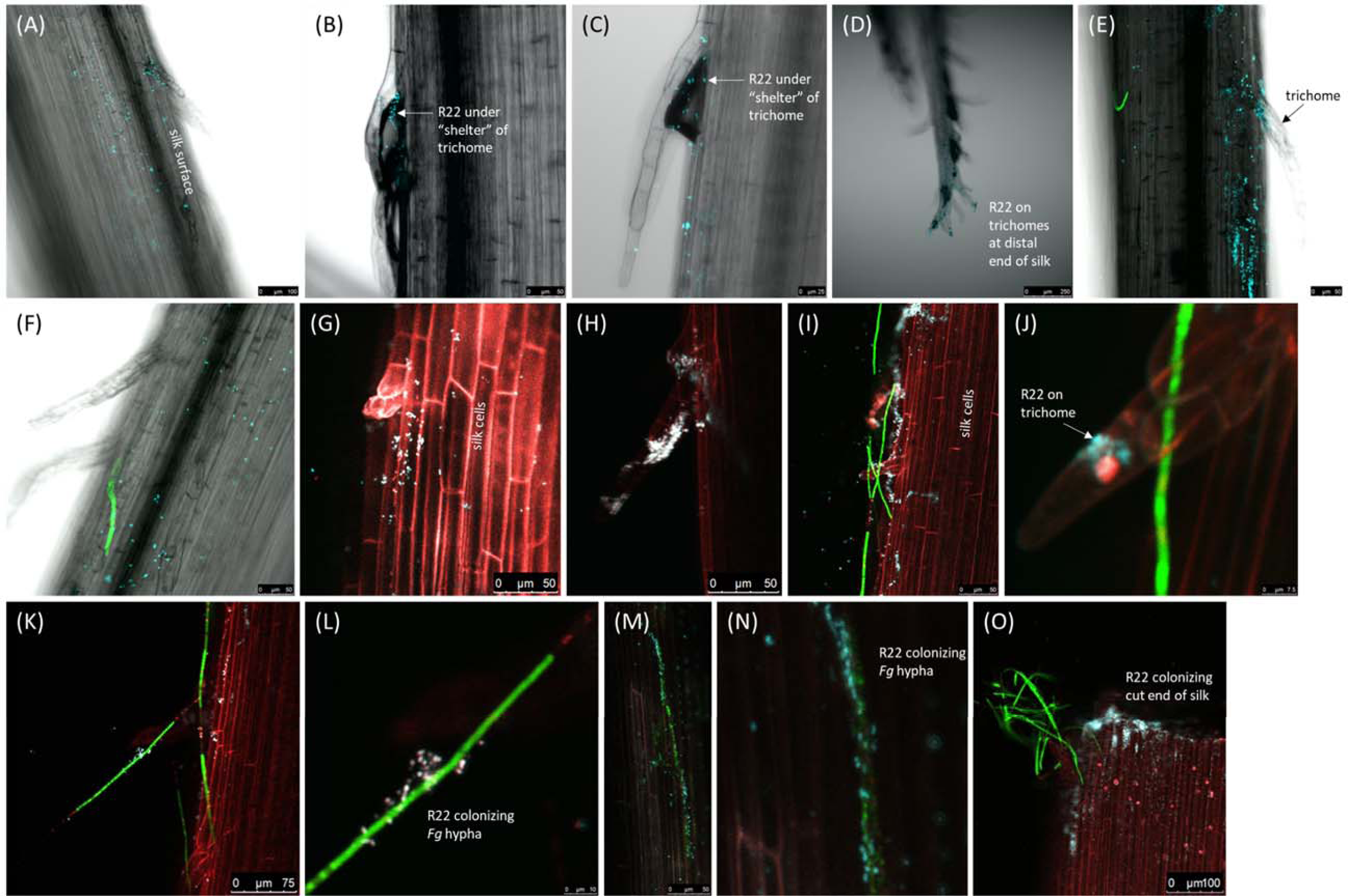
Confocal microscopy interactions between bacterial isolate DsRed-tagged *Pantoea ananatis* R22, previously isolated from fertilization-stage maize silks, and GFP-tagged *Fusarium graminearum* which enters maize from the environment at silk tips. Shown here are *Fg* colonization (*Fg*) and R22 colonization (digitally altered to cyan to distinguish from red plant propidium iodide staining) on the surface of maize silks (dark grey visible by compound light, and stained red with propidium iodide for confocal), trichomes, and the inside of cut silks. Isolate R22 did not match any identified V4-MiSeq OTUs, nonetheless had impressive interactions with silks and *Fg*. Two variations of *Method 1* were used. The first variation is shown in panels **(A-F),** where silks connected to a cob were placed on a disc of water agar in a Petri dish, and DsRed-tagged R22 in Tet-LB was applied to the silks on the water agar, and *Fg* was applied 1 cm from the water agar, and the silks were incubated for 2 d. Images of DsRed-tagged R22 and GFP-*Fg* captured with a confocal laser were overlayed with silks captured with a compound light microscope. The results were as follows: **(A)** R22 colonizing the silk surface, including a trichome. **(B)** R22 under silk trichomes, possibly with an air bubble (dark space under trichome). **(C-D)** R22 colonizing trichomes on the distal end of a silk. **(E)** R22 colonizing the silk surface and a trichome. **(F)** R22 colonizing the silk surface near *Fg* and silk trichomes. The second variation of Method 1 is shown in panels **(G-O)** where silks were prepared using the same methods, except for the application GFP-*Fg*, which occurred after 1 d of incubation, which was followed by an additional 2 d (3 d total incubation), and confocal fluorescence was used for all components. The results that are shown were purely confocal overlays: **(G)** R22 colonizing the silk surface. **(H)** R22 densely colonizing a silk trichome. **(I)** R22 on the silk surface, near transition sites where *Fg* was turning from green (live) to red (dead), also on trichomes. **(J)** R22 on a trichome, with an *Fg* hypha. **(K)** R22 directly colonizing an *Fg* hypha on the surface of a silk, near transition sites where *Fg* was turning from green (live) to red (dead). **(L)** A magnified version of the hypha shown in K, with R22 colonizing. **(M)** R22 coating a long stretch of *Fg* hypha. **(N)** A magnified version of the R22-coated hypha shown in M. **(O)** R22 entering and colonizing the inside of a silk at a cut end, near *Fg*.

**Figure 3.**
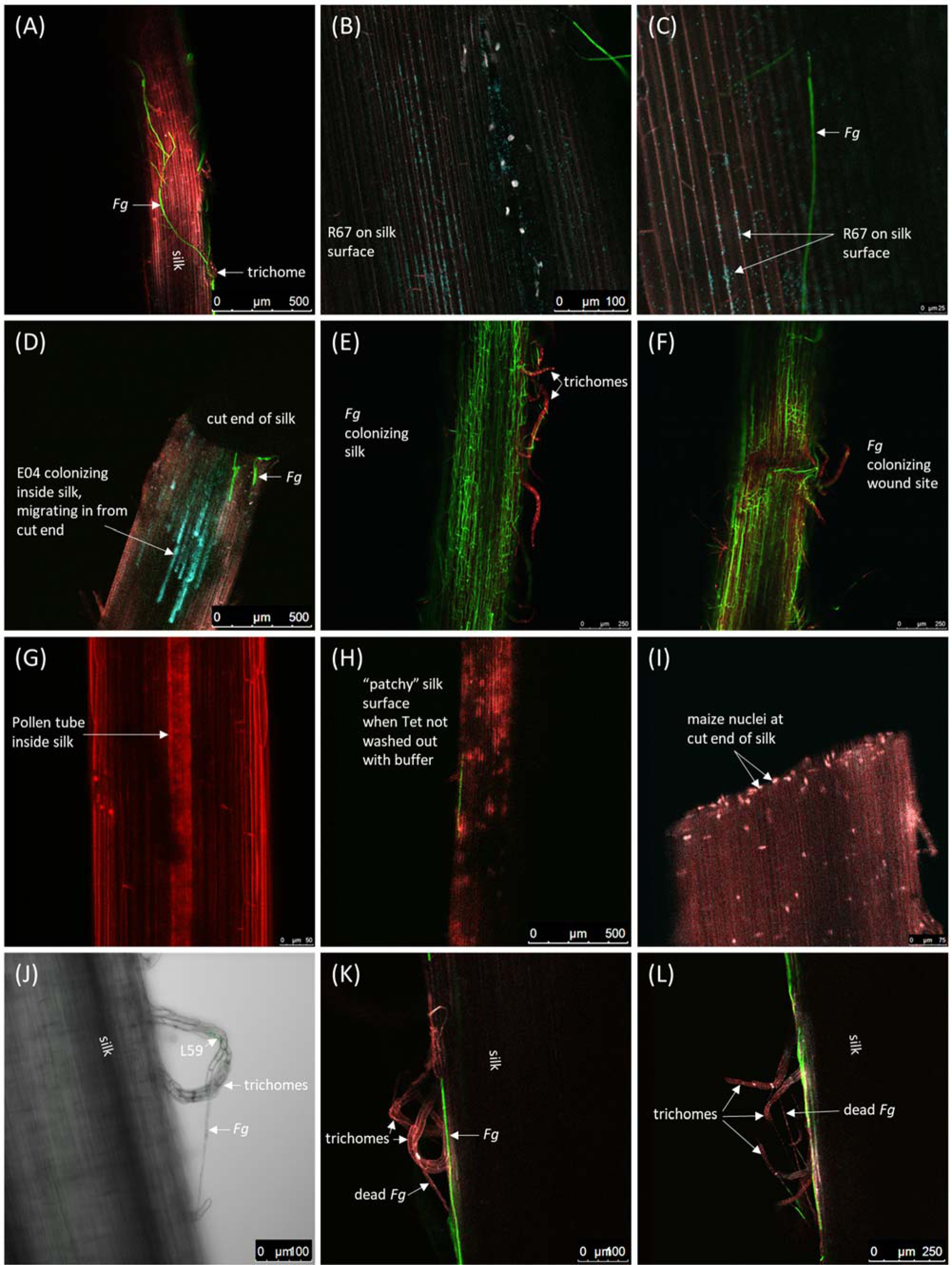
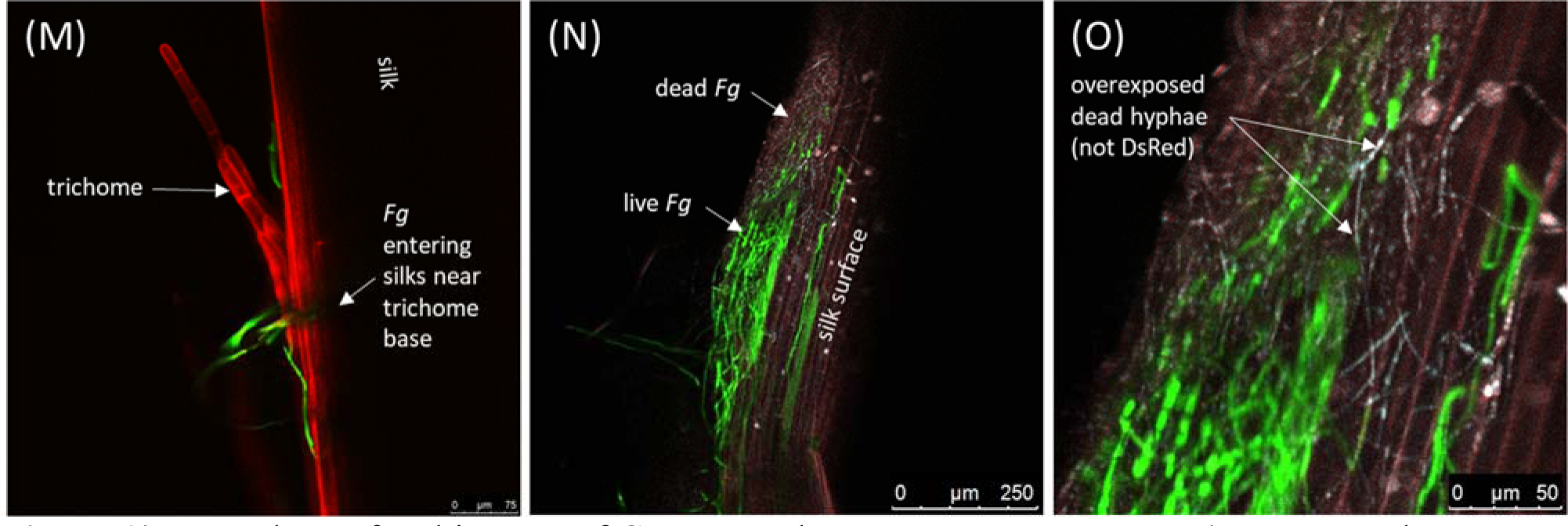
Example confocal images of GFP-tagged *Fusarium graminearum* (*Fg*, green when live, red when dead) and three DsRed-tagged bacterial endophytes (cyan) interacting on maize silks (red, stained with propidium iodide). These endophytes were previously isolated from fertilization-stage maize silks and include: R67 (*Pseudomonas*), E04 (*Pantoea*), and L59 (*Delftia*). **(A)** *Fg* colonizing the surface near the base of trichomes. Cut silks were placed in small water agar plate, and treatments of pH 7 Tris buffer and GFP-*Fg* that had been washed with the buffer 3X were applied to the silks, and the silks were incubated for 3 d. **(B-C)** DsRed-tagged R67 and *Fg* hyphae on silk surface. DsRed-tagged R67 in Tet-LB was applied to silks that were pre-cut, then incubated for 1 d and treated with GFP-*Fg*, and then incubated for an additional 2 d (3 d total). **(D)** E04 demonstrating endophytic potential, colonizing the inside of a maize silk, migrating inward from the cut end. Cut silks were placed on a small water agar plate, and both DsRed-tagged E04 and GFP-*Fg* were washed with pH 7 Tris buffer 3X and applied to the silks, and the silks were incubated for 3 d. **(E)** *Fg* colonizing a silk. **(F)** *Fg* colonizing a wound site on a silk. In (E-F), cut silks were placed in a small water agar plate, then pH 7 Tris buffer 3X and GFP-*Fg* were applied and the silks were incubated for 3 d. **(G)** A pollen tube visible within a silk. Silks connected to a cob were placed on a disc of water agar in a Petri dish, and DsRed-tagged R67 (not visible here) in Tet-LB was applied to the silks on the WA, incubated for 1 d, and then GFP-*Fg* was applied 1 cm from the WA, and the silks were incubated for an additional 2 d (3 d total). **(H)** A “patchy” appearance on the surface of a maize silk, commonly observed when tetracycline was not washed out of the bacterial treatments. Silks were prepared in the same manner as (G), but sterile Tet-LB was applied instead of DsRed-R67. **(I)** The cut end of a maize silk. Maize nuclei have migrated toward the cut end. Silks connected to a cob were placed on a disc of water agar in a Petri dish, and DsRed-tagged R67 (not visible here) in Tet-LB with 0.125% v/v Agral 90 (registration #11809, Syngenta Canada, Guelph, Canada) was applied to silks, then incubated for 1 d and treated with GFP-*Fg*, and then incubated for an additional 2 d (3 d total). **(J-L)** *Fg* hyphae appearing to “tether” maize trichomes to the main body of the silks. In (J), GFP-tagged L59 captured with a confocal laser, overlayed with untagged *Fg* and unstained silks captured with a compound light microscope. Silks connected to a cob were placed on a disc of water agar in a Petri dish, and GFP-tagged L59 was washed with pH 7 Tris buffer 3X and applied to the silks on the water agar disc, incubated for 1 d, and then *Fg* was applied 1 cm from the water agar, and the silks were incubated for an additional 4 d (5 d total). In (K-L), cut silks were placed in small water agar plate, and DsRed-tagged E04 (not visible here) was washed with pH 7 Tris buffer 3X and applied to the silks on the water agar, GFP-*Fg* was also applied, and the silks were incubated for 4 d. **(M)** *Fg* colonizing the surface of silks and penetrating near the base of trichomes. Silks connected to a cob were placed on a disc of WA in a Petri dish, and either a Tet-LB control or a DsRed-tagged bacterial treatment (not pictured) was applied to the silks on the WA, and GFP-*Fg* was applied at the distal end of the silks, and the silks were incubated for 2 d. **(N)** Live and dead *Fg* hyphae on the surface of a silk. **(O)** A magnified version of the hypha shown in panel Q; when dead *Fg* hyphae are over-exposed with the laser used for perceiving DsRed, they show some fluorescence (faint blue) although there is no DsRed present. In (N-O), cut silks were placed in small, empty Petri dish, and pH 7 Tris buffer 3X and GFP-*Fg* were applied, and the silks were incubated for 3 d.

### Observations of Method 1 *(silks attached to the cob)* with isolate R22, *Pantoea ananatis,* with *Fg* on maize silks

Method 1 allowed successful observation of featured anti-*Fg* bacterial isolate R22 colonizing the surface of maize silks, trichomes, *Fg* hyphae, and the inside of cut silks (Figure 2). Isolate R22 colonized maize silks, particularly well under trichomes (Figure 2B, C, E, H) and at silk tips (Figure 2D), and was observed colonizing the inside of silk after entering through a cut end (Figure 2O). Isolate R22 also densely colonized both live *Fg* hyphae (Figure 2M, N), and hyphae that was in the process of dying (transitioning from green to red; Figure 2K, L).

### Observations of Method 2 *(silks detached from the cob),* unique observations, and protocol modifications

Example images including silk-derived bacterial isolates R67 (*Pseudomonas*) and E04 (*Pantoea*) are included (Figure 3). Method 2 enabled successful observation of dense colonization of silk tissue by bacterial isolates and *Fg*, and the modified protocol showed interactions near and inside of silk wound sites. Example images show microbes on the silk surface, and internal colonization of silks via wound sites. Additionally, observations included a pollen tube visible inside of a silk, a “patchy” appearance on silks where Tetracycline had not been washed out of the treatment (Figure 3H), what are likely maize nuclei having migrated to the cut end of a silk (Figure 3I), and what appears to be “tethering” of trichomes by *Fg* (Figure 3J-L).

## DISCUSSION

Here, two novel pre-imaging protocols were presented to enable confocal fluorescence microscopy of maize silks. Using these methods, interactions between candidate silk-derived bacterial endophytes and the fungal pathogen *Fusarium graminearum* (*Fg*) that enters maize via silks, were successfully imaged on living maize silks under controlled conditions. These methods are biologically relevant, experimentally accessible, flexible to allow imaging diverse members of the microbiome, allow many replicates to be made simultaneously, and are practical for containment and biosafety purposes.

### Isolate R22, *Pantoea ananatis,* with *Fg* on maize silks

Of the four bacterial strains imaged, R22 (*Pantoea ananatis*) was particularly interesting for its ability to suppress *Fg* growth and colonization traits on silks. As a species, *Pantoea ananatis* includes some strains that are pathogenic to maize, and some strains that are maize endophytes (Sheibani-Tezerji et al., 2015). A strain of *Pantoea ananatis* was recently shown to reduce the *Fg*-produced mycotoxin deoxynivalenol in wheat (Deroo et al., 2022).

Method 1 (silks attached to the cob) was successfully used to image bacterial isolate R22 with *Fg* on maize silks. Isolate R22 did not match any identified V4-MiSeq OTUs in previous microbiome analyses (Thompson et al., n.d.-b), but was still a relevant anti-*Fg* strain *in vitro*.

However, there were strict thresholds in the bioinformatic analyses of the MiSeq results (Khalaf et al., 2021), so it is reasonable to expect that many strains of anti-fungal bacteria (including R22) would have been missed in the previous MiSeq microbiome analyses and not assigned a V4-MiSeq OTU.

Here, R22 was observed *in planta*, where it grew colonized under the “shelter” of trichomes, at the tips of silk, and entered into silk via a cut end (Figure 2). The fact that R22 entered the cut end of the silk in this instance was particularly impressive, because the silks had been uncut and attached to the cob for incubation, and were only cut for staining on the day of microscopy. This invasion of the silk only occurred within ∼1-2 hours of cutting and staining the silk.

Isolate R22 was also observed densely colonizing *Fg* hyphae, including near transition points where *Fg* was dying (turning from green to red). Isolate R22 contained genes for phenazine biosynthesis, acetoin biosynthesis, and colicin V biosynthesis (Thompson, 2023).

These images provide evidence that anti-*Fg* bacteria can be observed interacting with *Fg* on maize silks via confocal microscopy; the methods presented here represent a useful protocol that may be used for a variety of applications, including furthering our understanding of microbiome dynamics during the reproductive stage.

### Unique observations

There were several additional noteworthy observations from this study. First, nuclei migrated to the cut end of the silks, possibly for local transcription of wound-response/repair genes (Figure 3I). Nuclei are known to move within plant cells in response to abiotic stimuli, developmental stage, and plant–microbe interactions (Griffis et al., 2014). Most applicable here, nuclei are known for actin-dependent migration to wound sites (Takagi et al., 2011). Second, it was very difficult to get clear images beyond the outer layer of silk cells, but some images of the bacteria and *Fg* invading from wound sites were obtained. A unique image of a pollen tube inside a silk was obtained (Figure 3G), perhaps due to high uptake of propidium iodide by the pollen tube, or natural autofluoresence. Third, aside from endophyte interactions, these methods can further the understanding of plant-pathogen interactions. For example, *Fg* was seen entering the silks at the base of trichomes (Figure 2F; Figure 3M), similar to observations made by Miller et al., (2007), and also tethering of trichomes in some cases (Figure 3J-L). Finally, when comparing treatments of bacteria that had been washed with buffer to unwashed bacteria applied in the Tet-LB liquid media, unusual “patchy” patterns appeared under the confocal microscope on the silks that had been treated with unwashed bacteria. These patterns were also seen with Tet-LB control treatments (Figure 3H). This could possibly be attributed to the tetracycline in the unwashed liquid cultures.

### Cautionary Notes

There are some notes of caution pertaining to the confocal microscopy methods shown here. First, maize cell nuclei were visible with the lasers intended for PI and DsRed, but could be distinguished from bacteria by their size, shape, and placement. Second, buffer control dead hyphae also picked up some of the laser intended for DsRed, but these spots are not tagged bacteria (Figure 3N & O). Finally, the *Fg* hyphae were explored for breakages that might have been caused by the anti-*Fg* bacteria. Although hyphae often appeared to be cut or severed in a single optical slide, it was imperative to scan throughout the Z profile to confirm whether a hypha was simply blocked by something, commonly a trichome (or dead hyphae) further up in the Z profile. Under standard settings, confocal microscopes only reveal one extremely thin layer of the sample, which can be deceptive. It should be recommended that a thick Z-stack and 3D movie be created to convincingly demonstrate any hyphal breakages.

### Suggested modifications

We propose two suggested modifications to these protocols. First, the water agar disc (Figure 1B) could be replaced with different media to provide different nutrients or pH conditions for physiological studies.

We also have a suggested alternative protocol (Method 3) which we term the “silk bridge”: This adapted protocol is a suggestion for testing particularly motile microbes and rapid colonizers (Figure 1K). In short, a short segment of silk can be suspended between two segments of water agar, and one microbe (e.g. anti-fungal bacterial endophyte) can be applied to one section of water agar while a silk pathogen (e.g. *Fg*) can be applied to the other section, and the microbes would then be allowed to grow across the elevated section of silk, with the intention of observing their meeting point. Preliminary experiments were conducted, and the specific suggested protocol can be as follows: First, the middle portion of water agar from a 60 mm diameter Petri dish was cut and removed, leaving sections of water agar on two sides.

Approximately 2.5 cm sections of fresh silks were cut and plated, so that they laid across the open space to form a “bridge.” Next, 300 µL of GFP-*Fg* was applied to one section of water agar, and 300 µL of Tris-HCl-washed bacterial liquid culture or 10 mM Tris-HCl control was applied to the other section of water agar (Figure 1K).

## Methodology Limitations

One challenge, specific to this study, is that *Fg* grows and dies on silks, and stained hyphae (red) will occur on control silks, so stained hyphae alone is not enough to presume bacterial anti-fungal activity. Additionally, the silks are not sterile; they are known to be inhabited by a plethora of microbes (Khalaf et al., 2021; Thompson et al., n.d.-a), and thus the existent microbiome is expected to be present in these experiments and may have an effect on the *Fg* in both treated and control silks. However, this is a realistic scenario, where bacteria would be interacting with many microbes in-field, so it is a practical experiment.

To mitigate plant-to-plant differences in the silk microbiome, silks from an individual cob can be used for both treatment and control plates. For example, when using silks attached to the cob, a researcher can use half of the cob for a microbial treatment, and the other half for a control. When using cut silks, silks can be divided from a single cob into multiple plates of treatments and controls.

One application of these protocols can be to observe breakage of fungal hyphae due to bacterial attack. As a word of caution, imagery as proof of breakage should require 3D imaging in the form of a Z-stack, because observing a single optical slide/sheet/plane of the sample can be misleading. This is unlike typical compound light microscopy where one essentially views all of the levels superimposed, akin to a maximum projection in confocal microscopy. Hyphae can bend through the plane, or trichomes can interfere with light transmission, often giving the impression of breakage, where upon further investigation of the 3D environment, one may discover that no breakage occurred.

### Future applications

These protocols are predicted to be useful for observing different pathogens and tissues, including time-series observations. Other silk-entering fungi include *Fusarium verticillioides* (Sacc.) Nirenberg, *F. proliferatum* (Matsush.) Nirenberg, *F. subglutinans* (Wollenw. & Reinking) (three causative agents of Fusarium ear rot), *Aspergillus flavus* Link, *A. parasiticus* Speare (two causative agents of Aspergillus ear rot), *Ustilago maydis* (DC) Corda (also known as *U. zeae*, *U. zeae mays*, causative agent of corn smut), and *Stenocarpella maydis* (Berk.) Sutton (previously known as *Diplodia maydis*, causative agent of Diplodia ear rot) (Thompson & Raizada, 2018). All fungi listed produce mycotoxins, except for *U. maydis*, which can co-infect with mycotoxigenic fungi (Abbas et al., 2015, 2017). The protocols presented here may be valuable in evaluating treatments to combat multiple pathogenic fungi simultaneously.

Endophytes interact in large microbial communities (Woo & Pepe, 2018), and these protocols are not limited to observing one endophyte at a time. Future studies should be conducted to observe multiple endophytes (with different fluorescent tags to distinguish them) in consortia with one another.

Expanding the knowledge base about beneficial microbe modes of action will contribute to the development of more biocontrol options or prebiotic treatments, which may in turn reduce the environmental impacts (and cost to farmers) caused by antimicrobials including fungicides.

## Supplemental Methods S1: Fluorescent tagging of bacteria

Fluorescent protein-tagged microbes were created by transforming anti-Fusarium microbes from the in vitro trial by preparing electrocompetent cells as previously described by Mousa, Shearer, et al. (2016) and briefly described below. Select successfully expressing strains were carried forward to confocal experiments:

### S1.1 Plasmid DNA isolation

The tetracycline (Tet) resistant plasmids pSW002-PpsbA-DsRed-Express2 (DsRed)(Addgene, 111257), and the kanamycin (Kan) resistant plasmid, pDSK-GFP (GFP), were isolated from LB liquid cultures using the Quantum Prep Plasmid Miniprep Kit (BIO-RAD, Cat# 732-6100) and quantified with a Qubit v1.2 fluorometer.

### S1.2 Anti-Fg competent cell preparation

Anti-*Fg* bacterial cultures were grown in 25 mL LB at 200 rpm, 30 °C until an OD600 reading of 0.2-0.3, at which point they were placed on ice for 15-20 min. The cultures were then centrifuged at 4000 x g for 15 min at 4 °C and the supernatant was discarded. Then the pellet was resuspended in cold ddH2O, centrifuged again with the same settings, and the supernatant was discarded; this process was completed 3 times. The pellet was then resuspended in 5 mL of ice-cold 10% glycerol, centrifuged again, and the supernatant was discarded. The pellet was finally resuspended in 100 µL 10% ice-cold glycerol and dispensed in 50 µL aliquots, frozen using liquid nitrogen and stored at -80 °C.

### S1.3 Electroporation/Transformation

Competent cells (50 µL) were combined with 2 µL plasmid and transferred into a pre-chilled electroporation 0.1 cm gap cuvette which was tapped to remove bubbles, and electroporated at 1.8 Kv in a MicroPulser (Bio-Rad, USA). Immediately, 1 mL of pre-warmed LB broth was added into the cuvette. Aliquots of 50 µL were plated onto pre-warmed Tet (10 µg/ml) or Kan (50 µg/ml) LB plates, for the Tet-resistant or Kan-resistant plasmids, respectively. Fluorescent colonies were identified under UV light, restreaked on Tet/Kan plates, grown in Tet/Kan liquid cultures, and preserved in -80 °C glycerol stocks.

### Supplemental Methods S2: Greenhouse growth conditions

Pails had holes drilled in the bottom and on the sides near the base for drainage, and were pre-sterilized with 1% Virkon, rinsed with water, and allowed to dry before being filled with Turface® and set on top of metal grates to allow thorough drainage. Bamboo stakes were also sterilized with Virkon and used to provide support for the plants as they grew. There was a lateral fan near the roof of the greenhouse, and an upright fan which was moved around the room on a regular basis to promote airflow and strong stalk growth. The plants were thinned once they were 1-1.5 feet tall, keeping the most uniform plants. Pests were monitored using sticky traps, and biological control insects were applied regularly.

Pails were fed by drip fertigation for 1 min every 10 min, with plain water for the first 3 days after transplanting, and then with a solution of 83 g/L 24-10-20 Drip Irrigation fertilizer (Plant-Prod, Canada) amended with MgSO4 (1.746 g/L) and injected at a rate of ∼1:200 until harvest. The Turface® was top-dressed with ∼1 tsp limestone per week.

The greenhouse conditions were 28 °C/20 °C using natural sunlight, supplemented with full spectrum lamps and GroLux bulbs (16 h:8 h light:dark photoperiod).

## Author Contributions

M.E.H.T. conceived of the study, performed all experiments, and wrote the manuscript. M.N.R. edited the manuscript and supervised the study.

## Acknowledgments

Scholarships were provided to M.E.H.T. by NSERC CGS-D, NSERC CGS- M, and a Food from Thought Research Assistantship. We thank Dylan Brettingham, Jacob Kaszas, and Sneha Sengupta for their assistance in growing greenhouse maize. We thank Dr. Kamal Khadka for growing field maize. We thank Dr. Rajagopal Subramaniam (AAFC, Ottawa) for the source of GFP-tagged *Fusarium graminearum.*

## References

1. Abbas, H. K., Shier, W. T., Plasencia, J., Weaver, M. A., Bellaloui, N., Kotowicz, J. K., Butler, A. M., Accinelli, C., de la Torre-Hernandez, M. E., & Zablotowicz, R. M. (2017). Mycotoxin contamination in corn smut (Ustilago maydis) galls in the field and in the commercial food products. Food Control, 71, 57–63. 10.1016/j.foodcont.2016.06.006

2. Abbas, H. K., Zablotowicz, R. M., Shier, W. T., Johnson, B. J., Phillips, N. A., Weaver, M. A., Abel, C. A., & Bruns, H. A. (2015). Aflatoxin and fumonisin in corn (Zea mays) infected by common smut Ustilago maydis. Plant Disease, 99(9), 1236–1240. 10.1094/PDIS-03-14-0234-RE

3. Aleklett, K., Hart, M., & Shade, A. (2014). The microbial ecology of flowers: An emerging frontier in phyllosphere research. Botany, 92(4), 253–266. 10.1139/CJB-2013-0166/ASSET/IMAGES/LARGE/CJB-2013-0166F1.JPEG

4. Cardinale, M. (2014). Scanning a microhabitat: Plant-microbe interactions revealed by confocal laser microscopy. Frontiers in Microbiology, 5, 94. 10.3389/fmicb.2014.00094

5. Cui, Z., Huntley, R. B., Zeng, Q., & Steven, B. (2020). Temporal and spatial dynamics in the apple flower microbiome in the presence of the phytopathogen Erwinia amylovora. The ISME Journal, 15(1), 318–329. 10.1038/s41396-020-00784-y

6. De Ruyck, K., De Boevre, M., Huybrechts, I., & De Saeger, S. (2015). Dietary mycotoxins, co-exposure, and carcinogenesis in humans: Short review. Mutation Research - Reviews in Mutation Research, 766, 32–41. 10.1016/j.mrrev.2015.07.003

7. Deroo, W., De Troyer, L., Dumoulin, F., De Saeger, S., De Boevre, M., Vandenabeele, S., De Gelder, L., & Audenaert, K. (2022). A novel in planta enrichment method employing Fusarium graminearum-infected wheat spikes to select for competitive biocontrol bacteria. Toxins, 14(3), 222. 10.3390/TOXINS14030222/S1

8. Diniz, G. de F. D., Cota, L. V., Figueiredo, J. E. F., Aguiar, F. M., da Silva, D. D., de Paula Lana, U. G., dos Santos, V. L., Marriel, I. E., & de Oliveira-Paiva, C. A. (2022). Antifungal activity of bacterial strains from maize silks against Fusarium verticillioides. Archives of Microbiology, 204, 89. 10.1007/S00203-021-02726-4/TABLES/4

9. Diniz, G. de F. D., Figueiredo, J. E. F., Lana, U. G. P., Marins, M. S., Silva, D. D., Cota, L. V., Marriel, I. E., & Oliveira-Paiva, C. A. (2022). Microorganisms from corn stigma with biocontrol potential of Fusarium verticillioides. Brazilian Journal of Biology = Revista Brasleira de Biologia, 82, e262567. 10.1590/1519-6984.262567

10. Griffis, A. H. N., Groves, N. R., Zhou, X., & Meier, I. (2014). Nuclei in motion: Movement and positioning of plant nuclei in development, signaling, symbiosis, and disease. Frontiers in Plant Science, 5, 129. 10.3389/fpls.2014.00129

11. Khalaf, E. M., Shrestha, A., Rinne, J., Lynch, M. D. J., Shearer, C. R., Limay-Rios, V., Reid, L. M., & Raizada, M. N. (2021). Transmitting silks of maize have a complex and dynamic microbiome. Scientific Reports, 11(1), 13215. 10.1038/s41598-021-92648-4

12. Kiesselbach, T. (1999). The Structure and Reproduction of Corn (D. Brown & S. Schaefer, Eds.; 50th Anniv). Cold Spring Harbour Laboratory Press.

13. McHughen, A. (2016). A critical assessment of regulatory triggers for products of biotechnology: Product vs. process. GM Crops & Food, 7(3–4), 125–158. 10.1080/21645698.2016.1228516

14. Miller, S. S., Chabot, D. M. P., Ouellet, T., Harris, L. J., & Fedak, G. (2004). Use of a Fusarium graminearum strain transformed with green fluorescent protein to study infection in wheat (Triticum aestivum). Canadian Journal of Plant Pathology, 26, 453–463. 10.1080/07060660409507165

15. Miller, S. S., Reid, L. M., & Harris, L. J. (2007). Colonization of maize silks by Fusarium graminearum, the causative organism of gibberella ear rot. Canadian Journal of Botany, 85(4), 369–376. 10.1139/B07-027

16. Mousa, W. K., Schwan, A., Davidson, J., Strange, P., Liu, H., Zhou, T., Auzanneau, F.-I., & Raizada, M. N. (2015). An endophytic fungus isolated from finger millet (Eleusine coracana) produces anti-fungal natural products. Frontiers in Microbiology, 6, 1157. 10.3389/fmicb.2015.01157

17. Mousa, W. K., Schwan, A., & Raizada, M. (2016). Characterization of antifungal natural products isolated from endophytic fungi of finger millet (Eleusine coracana). Molecules, 21(12), 1171. 10.3390/molecules21091171

18. Mousa, W. K., Shearer, C., Limay-Rios, V., Ettinger, C. L., Eisen, J. A., & Raizada, M. N. (2016). Root-hair endophyte stacking in finger millet creates a physicochemical barrier to trap the fungal pathogen Fusarium graminearum. Nature Microbiology, 1(12), 16167. 10.1038/nmicrobiol.2016.167

19. Mousa, W. K., Shearer, C. R., Limay-Rios, V., Zhou, T., & Raizada, M. N. (2015). Bacterial endophytes from wild maize suppress Fusarium graminearum in modern maize and inhibit mycotoxin accumulation. Frontiers in Plant Science, 6, 805. 10.3389/fpls.2015.00805

20. Ngugi, H. K., & Scherm, H. (2006). Biology of flower-infecting fungi. Annual Review of Phytopathology, 44, 261–282. 10.1146/ANNUREV.PHYTO.44.070505.143405

21. Oldenburg, E., Höppner, F., Ellner, F., & Weinert, J. (2017). Fusarium diseases of maize associated with mycotoxin contamination of agricultural products intended to be used for food and feed. Mycotoxin Research, 33(3), 167–182. 10.1007/s12550-017-0277-y

22. Palmieri, D., Vitale, S., Lima, G., Di Pietro, A., & Turrà, D. (2020). A bacterial endophyte exploits chemotropism of a fungal pathogen for plant colonization. Nature Communications, 11, 5264. 10.1038/s41467-020-18994-5

23. Sankaranarayanan, A., Amaresan, N., & Dwivedi, M. K. (2023). Endophytic microbes: Isolation, identification, and bioactive potentials (A. Sankaranarayanan, N. Amaresan, & M. K. Dwivedi, Eds.; 1st ed.). Springer Science and Business Media, LLC. https://link.springer.com/10.1007/978-1-0716-2827-0

24. Sauter, M. (2009). A guided tour: Pollen tube orientation in flowering plants. Chinese Science Bulletin, 54(14), 2376–2382. 10.1007/s11434-009-0329-6

25. Schaafsma, A. W., Nicol, R. W., & Reid, L. M. (1997). Evaluating commercial maize hybrids for resistance to gibberella ear rot. European Journal of Plant Pathology, 103(8), 737–746. 10.1023/A:1008629629069

26. Sheibani-Tezerji, R., Naveed, M., Jehl, M. A., Sessitsch, A., Rattei, T., & Mitter, B. (2015). The genomes of closely related Pantoea ananatis maize seed endophytes having different effects on the host plant differ in secretion system genes and mobile genetic elements. Frontiers in Microbiology, 6, 440. 10.3389/FMICB.2015.00440/ABSTRACT

27. Takagi, S., Islam, M. S., & Iwabuchi, K. (2011). Dynamic behavior of double-membrane-bounded organelles in plant cells. In International Review of Cell and Molecular Biology (Vol. 286, Issue C, pp. 181–222). Elsevier Inc. 10.1016/B978-0-12-385859-7.00004-5

28. Thompson, M. E. H., & Raizada, M. N. (2018). Fungal pathogens of maize gaining free passage along the silk road. Pathogens, 7(4), 81. 10.3390/pathogens7040081

29. Thompson, M.E.H. Discovery and testing of pollinated maize silk-associated microbes including microbiome assisted selection of biocontrol agents against Fusarium graminearum, submitted PhD Thesis, University of Guelph, Canada, 2023.

30. Thompson, M. E. H., Shrestha, A., Rinne, J., Limay-Rios, V., Reid, L., & Raizada, M. N. (n.d.- a). Systematic Culturing and Functional Gene Mining of the Microbiome of Fertilization- Stage Silks (Style Tissue) of North American Maize Reveals Diversity and Possible Survival Traits to Promote Host Reproduction and Tolerate the Pollen and Silk Niches. Under Review.

31. Thompson, M. E. H., Shrestha, A., Rinne, J., Limay-Rios, V., Reid, L., & Raizada, M. N. (n.d.- b). The cultured microbiome of pollinated maize silks shifts after infection with Fusarium graminearum and varies by distance from the site of pathogen inoculation. *Under Review*.

32. War Nongkhlaw, F., & Joshi, S. (2017). Microscopic study on colonization and antimicrobial property of endophytic bacteria associated with ethnomedicinal plants of Meghalaya. Journal of Microscopy and Ultrastructure, 5(3), 132. 10.1016/j.jmau.2016.09.002

33. Woo, S. L., & Pepe, O. (2018). Microbial consortia: Promising probiotics as plant biostimulants for sustainable agriculture. Frontiers in Plant Science, 9, 1801. 10.3389/FPLS.2018.01801/BIBTEX

